# Decoding Protein Aggregation through Computational Approach: Identification and Scoring of Aggregation-Prone Regions in Protein Sequences

**DOI:** 10.1101/2024.06.11.598423

**Authors:** Rahul Kaushik, Thomas Launey

## Abstract

Protein aggregation is a critical phenomenon associated with numerous neurodegenerative and systemic diseases. Understanding the propensity of proteins to aggregate is essential for unraveling the molecular basis of these disorders and for design and engineering of novel proteins or modulating the activity/stability of enzymatic proteins. Here, we present APR-Score, a novel machine-learning based computational method designed to identify aggregation-prone regions within protein sequences. ARP-Score leverages a combination of sequence-based features to predict regions of proteins that are prone to aggregate. The APR-Score harnessed the information ingrained in the compiled sequence and structural features to provide state-of-the-art accuracy. The APR-Score is assessed by conducting rigorous cross-validation experiments on the training dataset and further validated on an independent test dataset. The APR-Score prediction models demonstrated robustness and reliability in discriminating aggregation-prone regions from non-aggregating ones on an independent dataset, achieving Mathew’s correlation coefficient (MCC) 0.81, precision 0.89, and F1-Score 0.91. The APR-Score offers a valuable tool for researchers investigating protein aggregation-related diseases, as it can expedite the identification of aggregation-prone regions, aiding in the development of targeted therapies and diagnostic tools. The computational protein design and engineering regimes can be facilitated through APR-Score based identification and screening of aggregation prone protein sequences.

## 1. Introduction

The tendency of proteins and peptides to form aggregates under different cellular environments is very common across the different forms of life and results into the subcategories either as amorphous aggregates or as insoluble ordered fibrous protein aggregates, characterized as amyloids (Hartl, 2017; Knowles, et al., 2014; Ono and Watanabe-Nakayama, 2021). The association of protein aggregation, majorly as amyloid fibrils, with different human diseases has been extensively established through various scientific studies. For instance, some of the human diseases are reported to have a crucial role of amyloid fibril formation like neurodegenerative diseases, viz. Alzheimer’s, Huntington’s and Parkinson (amyloid-β, tau or α-synuclein in the brain), type II diabetes (amylin in pancreas), systematic amyloidosis (transthyretin in different organs), cataract (α-crystallin in the eyes), and aggregation of p53 and Axin proteins in cancer cells (Hartl, 2017; Knowles, et al., 2014; Ono and Watanabe-Nakayama, 2021; Peng, et al., 2020; Sigurdson, et al., 2019). Furthermore, the induction of adverse immune reactions resulting from unintended protein aggregation poses serious challenges to formulate therapeutic proteins and peptides, and success of various drug discovery regimes. Better understanding of the protein aggregation signatures by identifying specific features in the protein sequences and their structural information could improve the end-product biopharmaceuticals (Hartl, 2017; Ono and Watanabe-Nakayama, 2021; Orlando, et al., 2020).

Different previous studies demonstrated the relevance of native state stability of protein in their aggregation propensities. The extent of native state stability of proteins that is contributed by aggregation prone regions is demonstrated to be higher as compared to other regions of the proteins. Interestingly, some proteins evolved in such a way that these governed the aggregation by excluding certain residues known for their role in aggregation. For instance, thermophilic proteins were found to be more efficient as compared to mesophilic proteins, in countering aggregation by means of either excluding specific residues or concealing them into the core of protein. The protein aggregate formation is facilitated by small aggregation susceptible polypeptides of usually 5-15 amino acid residues, generally termed as aggregation prone regions (Ebo, et al., 2020; Houben, et al., 2022; Meric, et al., 2021; Vendruscolo and Fuxreiter, 2023) . A variation in the conditions or mutations initiates their exposure and thus assembly into inter-molecular beta sheets, resulting in the nucleation of aggregation of proteins (Gallardo, et al., 2020; Langenberg, et al., 2020; Seuma, et al., 2022). Several state-of-the-art approaches have been developed recently for identifying and scoring the aggregation prone regions in proteins. The susceptibility of α-helices to form β-sheets for certain segments of protein sequences was utilized for searching aggregates forming proteins. The detailed studies of these conformational switches from helices to sheets laid the foundation of various aggregation prediction approaches (Eshari, et al., 2023; Smith and Shell, 2017). The studies related to the impact of mutations on peptides and proteins aggregations helped the computational characterization and identification of aggregation propensities of polypeptides (Fernandez-Escamilla, et al., 2004; Smith and Shell, 2017). Various mathematical and machine learning approaches, including position-specific scoring matrices approach, Bayesian classifier and weighted decision tree approach, statistical mechanics algorithm, etc., are being implemented on the knowledge derived from protein sequence and structural properties to identify and score aggregation prone regions in proteins. Some of these properties include the extent of hydrophobicity and residue composition of small segments, residue pair preferences, β-strand propensity, and solvent accessibility/burial of amino acid residues in protein structures (Beerten, et al., 2015; Co, et al., 2022; Conchillo-Solé, et al., 2007;Fernandez-Escamilla, et al., 2004; Garbuzynskiy, et al., 2010; O’Donnell, et al., 2011; Smith and Shell, 2017; Walsh, et al., 2014; Zhang, et al., 2007).

Despite the development of several methods for identifying the aggregation prone regions, the desired accuracy and consistency is yet to be achieved on a diverse dataset. Considering the essence of the field, here we propose a set of highly reliable machine learning based methods, named APR-Score, for identification and scoring of aggregation prone regions in proteins that could deliver some novel insights into the self-assembly problems. The methodology integrates the features of amino acids accounting for their physico-chemical, sequence and structural properties along with their aggregation propensities derived from experimental studies and utilize them using different machine learning approaches to achieve improved predictions. The proposed methodology is thoroughly validated and benchmarked on a dataset of experimentally characterized aggregate forming proteins.

## 2. Methodology

### 2.1 Initial Data Compilation

An initial dataset of experimentally characterized aggregation and amyloid forming polypeptides was compiled from some well-known databases of sequence/structural information of aggregate/amyloid-forming peptides, viz. WALTZ-DB 2.0 (Louros, et al., 2020), CPAD 2.0 (Rawat, et al., 2020), AmyPro (Varadi, et al., 2018), and AmyloGraph (Burdukiewicz, et al., 2023). The polypeptides with redundant information in terms of sequence, position in the protein sequences, and experimental characterization were filtered out to avoid any bias. Notably, a major fraction of these experimentally characterized polypeptides was composed of hexapeptides that was used (partially or completely) as a reliable source of information for devising principles for identifying and scoring aggregation prone regions in proteins. Also, several experimental studies demonstrated the hexapeptides as the very critical units involved in initiating the aggregation or for the nucleation of amyloid formation (Beerten, et al., 2015; Teng and Eisenberg, 2009; Zbilut, et al., 2006). Interestingly, two cases were observed among the experimentally characterized hexapeptides where.the same hexapeptide was characterized as both non-aggregating and aggregating in the same protein at same residue positions (hexapeptides NNQQNY at 8^th^ residue position and KNFNYN at 102^nd^ residue position of UniProt Id: P05453). Likewise, there was one case of experimentally characterized hexapeptide as amyloid (Sequence: VTQVGF) in same protein (UniProt Id: P28307) at two different residues positions (at 45^th^ residue and at 137^th^ residue). These three hexapeptides were not included in the selected dataset of 1101 hexapeptides and were studies separately as described in the results section. From the compiled non-redundant dataset, a refined dataset of 1101 mutually exclusive hexapeptides having defined UniProt Identifiers was extracted and used further in this study. Among the 1101 hexapeptides, 341 hexapeptides were labelled as aggregate forming and 760 hexapeptides were non-aggregates forming. To avoid any bias in developing the proposed prediction model, the non-aggregate forming hexapeptides were randomly under-sampled to 341 hexapeptides, leading to a non-redundant dataset of 682 hexapeptides with their experimental characterization as amyloid/aggregate or non-amyloid/non-aggregates forming. This dataset was further portioned into Training dataset (n = 546) and Test dataset (n = 136). Notably, both the Training (n_aggregating_ = 273, n_non-aggregating_ = 273) and Test (n_aggregating_ = 68, n_non-aggregating_ = 68) datasets had equally distributed aggregate and non-aggregates forming hexapeptides.

### 2.2 Sequence and Structural Features Compilation

For developing the aggregation prone region score (APR-Score) predictor, the parameters that are experimentally reported to be significant for aggregate or amyloid formation were accounted. A total of nine parameters that could be derived from protein sequence information of the Training Dataset were selected as the input for training the initial neural network-based machine learning models. The sequence-based features were derived from hydrophobicity score, packing density, secondary structural propensities, residues depth propensities, mean polarity, partition energy, linker index, and Shannon’s entropy. The values of the selected sequence and structure-based features for individual amino acids were adopted from previous experimental studies and provided in supplementary information (Table S1).

#### 2.2.1 Features Derived from Hydrophobicity Scores

Hydrophobicity is reported among the crucial deriving forces for aggregate formation in several experimental studies as hydrophobic residues lead to higher aggregation tendencies as compared to polar residues. Most of the aggregation prone regions in protein sequences are reported to be very rich in hydrophobic residues (Breydo, et al., 2015; Maurya, et al., 2022; Ptak-Kaczor, et al., 2021; Xiao, et al., 2022). For developing APR-Score, we adopted Naderi-Manesh hydrophobicity scale (Naderi-Manesh, et al., 2001) and normalized the scale for on a scale of -1 to 1, where 1 represented the most hydrophobic residue and vice versa. The normalized hydrophobicity scale was used to derive different hydrophobicity-based features. Each individual residue in a protein sequence was designated as hydrophobic or non-hydrophobic based on its own hydrophobicity score along with the hydrophobicity score of its N-terminal and C-terminal neighboring residues. If the mean hydrophobicity score of three consecutive residues counted greater than zero, the middle residue was assigned as hydrophobic residue and vice-versa. For the first residue (without N-terminal neighbor) and the last residue (without C-terminal residue) of the protein sequence, only the available neighboring residue was accounted for the assignment. The following features were derived at hexapeptide level (using a 6 amino acid residues sliding window) from the residue assignment and the hydrophobicity scores, (i) effective ratio of hydrophobic and non-hydrophobic residues ((number of hydrophobic residues - number of non-hydrophobic residues)/ (number of hydrophobic residues + number of non-hydrophobic residues)), (ii) mean hydrophobicity score of hydrophobic residues, (iii) mean hydrophobicity score of non-hydrophobic residues, (iv) overall hydrophobicity score of hexapeptide irrespective of individual residue assignment. These four features were calculated for the hexapeptides compiled in section 2.1, using the corresponding scores for individual amino acid residues for hydrophobicity and mean polarity (Table S1).

The presence of polar amino acid residues imparts an enhancement in protein solubility and thus affects the aggregate formation significantly as reported in previous studies (Cukalevski, et al., 2012; Madhusudan Makwana and Mahalakshmi, 2015; Skibiszewska, et al., 2020; Stanković, et al., 2020). Considering this, the mean polarity of individual amino acid residues was also accounted asan important parameter for aggregation prone regions in the protein sequences. A similar set of four parameters was derived from the normalized mean polarity scores of individual residues, resulting into a set of eight parameters that account for polar and non-polar amino acid residues at hexapeptide level. In total, eight parameters derived from hydrophobicity and mean polarity scores were implemented in the development of the proposed APR-Score predictor.

#### 2.2.2 Features Derived from Packing Densities of Residues

The packing densities of amino acid residues reflect their preferential environment of neighboring residues in the 3-dimensional space. Depending on the extent of packing, amino acid residues are observed to be involved in aggregate initiation and formation (Fang, et al., 2013; Guo and Akhremitchev, 2006; Housmans, et al., 2023; Santos, et al., 2020). Considering the experimentally demonstrated role of packing densities of residues in aggregate formation, it was commissioned as one of the key features for devising the proposed APR-Score predictor. The reference values for packing densities of individual residues were adopted from experimentally solved proteins (clustered at 25% sequence identity) with 8Å contact radius (Galzitskaya, et al., 2006). The packing densities for individual residues are normalized to a scale ranging from -1 to 1 (Table S1), where -1 represented most loosely packed residue and 1 represented the most densely packed residue. The individual residues in the input sequence were assigned as densely packed or loosely packed based on their packing density scores along with the packing densities of its N-terminal and C-terminal neighboring residues. A positive mean packing density score of three consecutive residues resulted in a densely packed middle residue and vice versa. Similar to hydrophobicity score, the first residue (without N-terminal neighbor) and the last residue (without C-terminal residue) of the protein sequence, only the available neighboring residues were accounted for the assignment. A set of following features was derived from the packing densities scores and residue assignment using a 6 residues (hexapeptide) sliding window, (i) effective ratio of densely packed and loosely packed residues ((number of densely packed residues - number of loosely packed residues)/ (number of densely packed residues + number of loosely residues)), (ii) mean packing density of densely packed residues, (iii) mean packing density of loosely packed residues, (iv) overall packing density of hexapeptide irrespective of individual residue assignment. These four features were calculated for all the hexapeptides as compiled in section 2.1.

#### 2.2.3 Linker Propensity Indices of Amino Acid Residues

The linker propensities of amino acid residues are inherently implemented for identifying and analyzing the domain-domain interactions among the same or different proteins. The linker propensity of a residue reflects its tendency to interact with other residues in its vicinity (Jones, et al., 2000; Zhang, et al., 2011). The residues with a high linker propensity tend to participate in various interactions and experimentally reported to impart the cooperativity when located in the aggregation prone regions of the long amyloidogenic protein sequences (Chiti and Dobson, 2006; Chiti and Dobson, 2009). For the development of APR-Score, the linker propensities of individual amino acid residues over a sliding window of six amino acid residues was used for detecting the potential aggregation prone regions. The linker propensity indices were normalized to a scale ranging from -1 to 1 (shown in Table S1), where 1 represented the most linked residue and vice versa. Each individual residue in a protein sequence was designated as linker residue or neutral residue based on its own linker propensity as well as the linker propensities of its N-terminal and C-terminal neighboring residues. If the mean linker propensity of three consecutive residues counted greater than zero, the middle residue was assigned as linker residue and vice-versa. The following features were derived at the hexapeptide level from the residue assignment and their linker propensities, (i) effective ratio of linker and neutral residues ((number of linker residues - number of neutral residues)/ (number of linker residues + number of neutral residues)), (ii) mean propensity of linker residues, (iii) mean propensity of neutral residues, (vi) overall propensity of the hexapeptide irrespective of individual residue assignment. These four features were calculated for all the hexapeptides compiled in section 2.1 and were used for developing the proposed APR-Score predictor.

#### 2.2.4 Partition Energies of Residues

The approximate spatial position of an amino acid residue in the protein structure can be predicted from its partition energy of transfer from polar to hydrophobic environment. Certain polypeptides with higher partition energy are known to play crucial roles in various protein-protein interactions (Houben, et al., 2022; Housmans, et al., 2023; Langenberg, et al., 2020; Shaytan, et al., 2009). The role of partition energies of the amino acid residues in protein-protein interaction projects partition energy as a potential candidate parameter to explore its role in aggregate formation. For developing the proposed APR-Score, the partition energies of amino acid residues (de Jong, et al., 2013; Guy, 1985) were used for identifying aggregation protein regions in the protein sequences. The partition energies of amino acids are normalized to scale ranging from -1 to 1 as shown in Table S1.

#### 2.2.5 Secondary Structural Compatibility (CSS) Score for β-sheet Propensity

It has been experimentally demonstrated that the residues with higher β-sheet propensities impart positive effect on aggregate formation in proteins (Chiti and Dobson, 2006; Chiti and Dobson, 2009; Co, et al., 2022; Cukalevski, et al., 2012; Ebo, et al., 2020; Hartl, 2017). It is also reported that sometimes the unstable α-helices assumes β-sheet confirmation as an effect of change in environment (O’Donnell, et al., 2011; Ono and Watanabe-Nakayama, 2021; Smith and Shell, 2017). The hexapeptide peptides that were having a similar α-helix and β-sheet forming propensities are more prone to switch from helical structure to β-sheet with small changes in their environment. Here, we utilized the β-sheet and α-helix propensities of the protein sequences at tripeptide level, adopted from CSS-Scores (Kaushik and Zhang, 2020; Kaushik and Zhang, 2022; Kaushik and Zhang, 2022). A reference library of CSS scores for all the tripeptides having all residues as β-sheet (e.g. VLV – EEE), normalized by the corresponding score for same tripeptide having all residues as helices (e.g. VLV – HHH) was compiled from previously reported CSS scores (Kaushik and Zhang, 2020). These normalized CSS-Scores at the hexapeptide level to account for the α-helix switching β-sheet propensities were calculated by applying sliding window approach of six amino acid residues and used as a parameter to identify the aggregation prone regions in the protein sequences. The scores close to unity were considered to be favoring α-helix to β-sheet switching, that could lead to protein aggregation.

#### 2.2.6 Shannon’s Entropy of Hexapeptides

The position specific propensities have been used for quantifying the conservation of residues at a given position in several sequence alignment related studies (Beerten, et al., 2015; Ebo, et al., 2020; Louros, et al., 2020; Maurer-Stroh, et al., 2010). The position specific propensities were used in the form of Shannon’s entropy, derived from the Theory of Information proposed by Claude Shannon for quantification of information. For devising APR-Score, we calculated the Shannon’s entropy corresponding to all individual residues at different positions in aggregate forming hexapeptides. First, the propensities of amino acid residues at position 1, 2, 3, 4, 5 and 6 in aggregate forming hexapeptide were calculated by normalizing the fraction of residue ‘x’ (x □ any standard amino acid residue) at position ‘i’ (i □ any position in hexapeptide, i.e. 1, 2, 3, 4, 5, and 6) by the total frequency of residue ‘x’ at any position. These propensities were used for calculating Shannon’s entropy (as shown in equation (1)), in the form of a 2D matrix of 20 rows (one for each amino acid) and 6 columns (one for each position in hexapeptide), as provided in supplementary Table S2

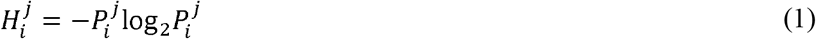

where, 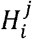 is Shannon’s entropy of an amino acid residue ‘x’ (x □ any standard amino acid residue) at position ‘i’ (1 ≤ i ≥ 6) in a hexapeptide.

These entropy values were calculated for the individual hexapeptide in the compiled dataset. In case of a protein sequence of interest, Shannon’s entropy for all possible hexapeptides was calculated by applying sliding window approach of six amino acid residues to identify the aggregation prone regions.

### 2.3 Prediction Model Development: Training and Testing

A comprehensive set of parameters/hyper-parameters (n = 22) accounting for six different features of hexapeptides of amino acid residues (as summarized in Table S3) in the initial dataset was used for selecting the most relevant parameters by implementing Principal Components Analysis (PCA). The selected parameters (n =18) were used for the development of the machine learning prediction model for identifying aggregation prone regions in protein sequences. The model was trained on the dataset of 546 hexapeptides that are experimentally characterized for their aggregation propensities and the developed model was evaluated on the Test dataset of 136 experimentally characterized hexapeptides. We developed five different machine learning models, viz., Random Forest (RF; 100 trees with replicable training and minimum subset not smaller than 10) based prediction model (APR-Score-RF), Support Vector Machine (SVM; with radial basis function implementing cost = 1.00, regression loss epsilon = 0.10, numerical tolerance = 0.001, and maximum iterations =100) based prediction model (APR-Score-SVM), Neural Network (NN; replicable training with 5 hidden layers implementing rectified linear unit activation function, and Adam solver function) based prediction model (APR-Score-NN), Gradient Boosting (GB; replicable training consisting 500 decision trees with a limited depth of 3 for each tree, and implementing learning rate 0.1) based prediction model (APR-Score-GB), and Logistic Regression (LR; with ridge regularization and balanced strength) based prediction model (APR-Score-LR). These prediction models were initially assessed based on their performance at 10-fold stratified cross validation and the assessment was further validated on the independent (never seen) Test Dataset.

### 2.4 Benchmarking of Prediction Models with other State of the Art Methods

The performance of different prediction models developed in the current study (APR-Score-RF, APR-Score-SVM, APR-Score-NN, APR-Score-GB, and APR-Score-LR) was benchmarked with the current state of the art methods for identifying aggregation prone regions (or amyloid formation initiating regions) in the protein sequences. These methods included TANGO (Fernandez-Escamilla, et al., 2004; Rousseau, et al., 2006), PASTA 2.0 (Walsh, et al., 2014), MetAmyl (Emily, et al., 2013), ANuPP (Prabakaran, et al., 2021), WALTZ (Beerten, et al., 2015; Louros, et al., 2020), and AGGRESCAN (Conchillo-Solé, et al., 2007; de Groot, et al., 2012). A brief description of these methods is provided in supplementary (Supplementary Note I). The performance of different prediction models developed in this study was benchmarked with the mentioned methods in terms of sensitivity, specificity, F1 score, precision, recall, and Mathew’s correlation coefficient (MCC).

## 3. Results and Discussion

### 3.1 Principal Components Analysis: Selection of Most Relevant Parameters

The compiled parameters were screened for their relevance in terms of imparting the cumulative variance among the features, to get rid of redundant parameters, and identifying the aggregation prone regions in the Training dataset. A Principal Components Analysis was performed to capture 99.9% cumulative variance from the compiled parameters. Based on the extent of Information Gain Ratio, and ANOVA, four parameters were identified as redundant and thus scrapped for developing the prediction models. This included the parameters derived from partition energy of amino acid residues (average partition energy, ratio of average partition energy of buried residues to the exposed residues, average partition energy of buried residues, and average partition energy of exposed residues). The corresponding values for Information Gain Ratio, and ANOVA in explaining the cumulative variance of the parameters are provided in supplementary (Table S3). The selected parameters (n =18) were further used for developing the prediction models by implementing different machine learning approaches. Among the selected parameters, the parameters derived from packing density, mean polarity and hydrophobicity, linker index, sequence-structure compatibility contributed to achieve a better performance of the proposed method. Further, the impact of individual set of parameters was studied to investigate the key deriving factors in identifying the aggregation prone regions in protein sequences.

### 3.2 Assessment of Individual Set of Parameters

Post-PCA, among the selected parameters (n =18), four parameters were derived from packing densities of amino acid residues, four parameters were derived from mean polarity scores of residues, a set of another four parameters was derived from linker indices of residues, four parameters accounted for hydrophobicity scores, one parameter accounted for sequence-structure compatibility scores, and one parameter considered amino acid residues position specificity ingrained through Shannon’s entropy. It is worth mentioning that the classification of an amino acid residues into densely or loosely packed, polar or non-polar, linked or neutral, hydrophobic or non-hydrophobic was derived from the cumulative score of its own score as well as its N-and C-terminal neighboring residues. Consideration of neighboring residues in categorizing the residues may have imparted an additional prediction accuracy in identifying aggregation prone regions. These six (packing density, mean polarity, linker index, hydrophobicity, CSS scores, and Shannon’s entropy) individual set of parameters were assessed independently in terms of area under receiver operating characteristic (ROC) curve (AUC), and recall for identifying aggregation prone regions through RF, SVM, NN, GB, and LR based machine learning prediction models through a stratified 10-fold cross validation. The results of the different prediction models for individual set of selected parameters are summarized in Table 1.

**Table 1.** A summary of area under receiver operating characteristic (ROC) curve and recall based assessment of prediction potential of individual set of parameters in identifying aggregation prone regions through different machine learning prediction models. The assessment scores are given in pairs where the first number represents the area under ROC curve, and the second number represents the recall.

| Individual Set of Parameters | Random Forest | Support Vector Machine | Neural Network | Gradient Boosting | Logistic Regression |
| --- | --- | --- | --- | --- | --- |
| Packing Density | 0.87; 0.82 | 0.84; 0.81 | 0.81; 0.78 | 0.86; 0.78 | <u>0.89</u>; 0.83 |
| Mean Polarity | <u>0.89</u>; 0.82 | 0.84; 0.79 | 0.85; 0.81 | 0.87; 0.79 | 0.88; 0.81 |
| Linker Index | 0.87; 0.80 | 0.85; 0.80 | 0.83; 0.80 | 0.83; 0.74 | <u>0.87</u>; 0.81 |
| Hydrophobicity | 0.83; 0.78 | 0.80; 0.72 | 0.80; 0.76 | 0.80; 0.73 | <u>0.85</u>; 0.77 |
| CSS Scores | 0.73; 0.66 | 0.55; 0.57 | <u>0.78</u>; 0.72 | 0.71; 0.61 | 0.76; 0.68 |
| Shannon's Entropy | <u>0.64</u>; 0.60 | 0.49; 0.48 | 0.63; 0.59 | 0.63; 0.59 | 0.63; 0.55 |

It is observed that the set of parameters derived from different features could achieve reasonable accuracy and recall metrics for the 10-fold cross validation. Notably, a simplistic approach implemented through Logistic Regression demonstrated equally good as compared to the other machine learning approaches accounted here. Interestingly, when analyzed in terms of area under ROC curve, the Logistic Regression based prediction model indicated better performance for three sets of parameters (packing density, linker index, and hydrophobicity) while the Random Forest based prediction model showed better statistics for mean polarity and Shannon’s entropy. The Neural Network based prediction model could extract better information in terms of area under ROC curve from sequence-structural compatibility scores. The training performance from the sets of features (n =6) showed a similar pattern in the case of the different statistics for Random Forest, Support Vector Machine, Neural Network, Gradient Boosting and Logistic Regression, as summarized in Table 1.

### 3.3 Cross Validation of Prediction Models for Identifying Aggregation Prone Regions

A stratified 10-fold cross validation on the training dataset was performed to evaluate the performance of different machine learning prediction models with the selected parameters (n =18). The rationale of this exercise was to identify the best performing prediction model for identifying the aggregation prone regions in protein sequences. A summary of the performances of different prediction models in 10-fold cross validation accounting for sensitivity, specificity, area under ROC curve (AUC), classification accuracy (CA), F1 score, precision, and recall metrics is provided in Table 2. The AUC metric is observed to be almost similar for all the prediction models, while the Random Forest based prediction model performed marginally better than other models in terms of classification accuracy, F1 score, and recall metrics. The ROC curves for individual prediction model are provided in supplementary Figure S1.

**Table 2.** A summary of performance evaluation of different prediction models in 10-fold cross validation accounting for sensitivity, specificity, area under ROC curve (AUC), classification accuracy (CA), F1 score, precision, and recall metrics.

| Prediction Model | Sensitivity | Specificity | AUC | CA | F1 Score | Precision | Recall |
| --- | --- | --- | --- | --- | --- | --- | --- |
| APR-Score-RF | <u>0.87</u> | 0.82 | <u>0.90</u> | <u>0.85</u> | <u>0.85</u> | <u>0.83</u> | <u>0.85</u> |
| APR-Score-SVM | 0.86 | 0.81 | 0.89 | 0.83 | 0.83 | 0.82 | 0.83 |
| APR-Score-NN | 0.82 | 0.79 | 0.88 | 0.81 | 0.81 | 0.80 | 0.81 |
| APR-Score-GB | 0.84 | <u>0.84</u> | <u>0.90</u> | 0.83 | 0.83 | 0.83 | 0.84 |
| APR-Score-LR | 0.84 | 0.81 | <u>0.90</u> | 0.82 | 0.82 | 0.81 | 0.82 |

Additionally, true positive rate, and false positive rate analysis at a threshold of 0.5 was performed for the different machine learning based prediction models and summarized in supplementary Table S4. The performance of different prediction models in identifying aggregation prone signatures ingrained at hexapeptide levels in protein sequences is depicted through a violin plot for bivariate analysis in Figure 1. The violin plot reflects the association between the predicted aggregation scores through different models and the category of non-aggregating (as 0) and aggregating (as 1) hexapeptides. The predicted aggregation scores are represented on x-axis and the aggregation / non-aggregation category is represented on y-axis. The plot illustrates the distribution of aggregation scores for aggregating and non-aggregating hexapeptides. It is observed that the distributions are mainly centered in the lower value regions of predicted aggregation scores for non-aggregating peptides while in the higher values regions of predicted aggregation scores for aggregating peptides.

**Figure 1.**
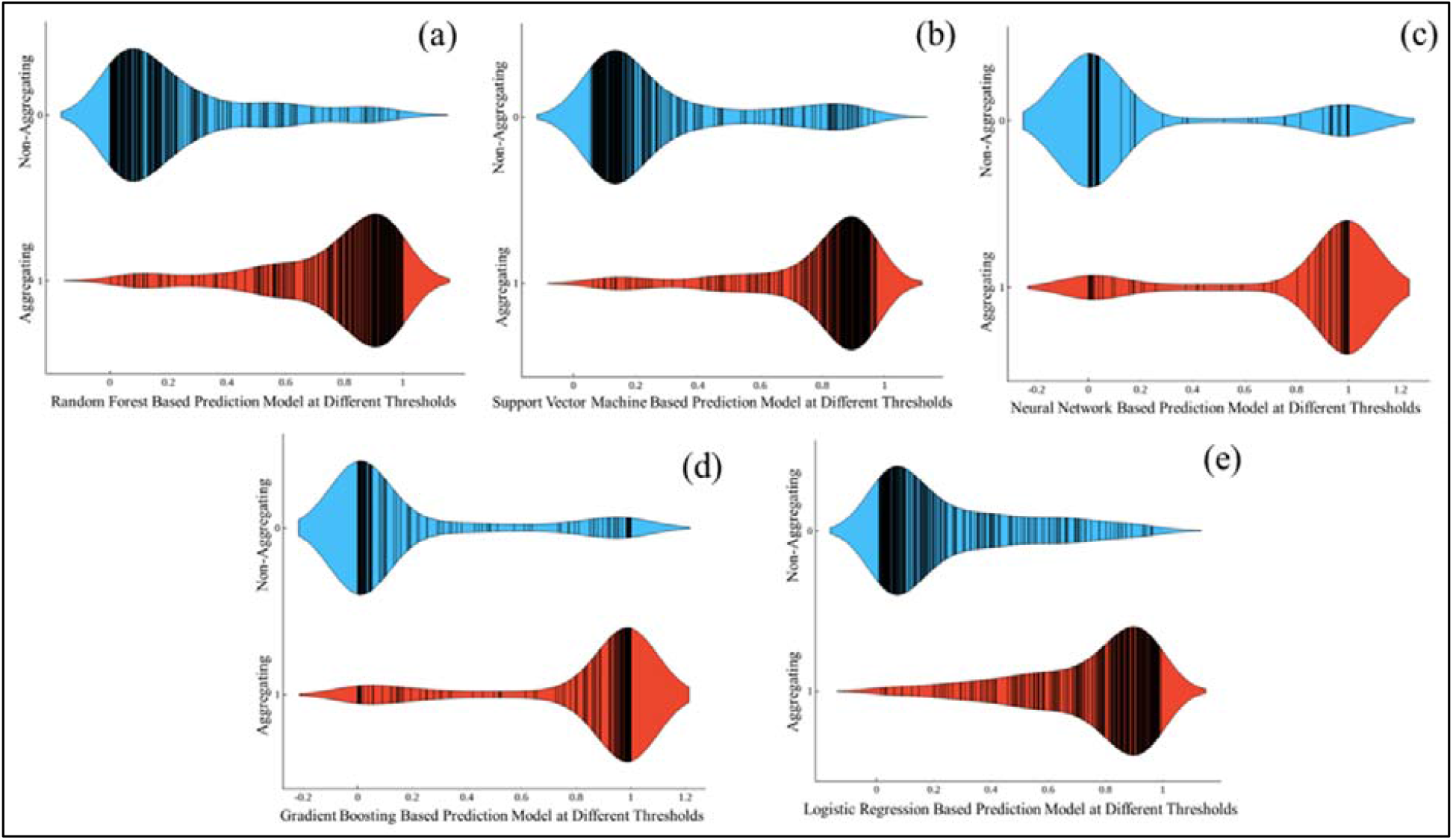
A violin plot depiction of the performances of different prediction models in identifying aggregation prone signatures ingrained at hexapeptide levels in protein sequences. The plot illustrates the distribution of predicted aggregation scores through different machine learning based prediction models, (a) APR-Score-RF, (b) APR-Score-SVM, (c) APR-Score-NN, (d) APR-Score-NN, and (e) APR-Score-LR for aggregating and non-aggregating peptides.

The possibility of achieving an improved performance by integrating the different prediction models was explored without having any gain over the best individual prediction model (APR-RF; Sensitivity = 0.87, AUC = 0.90, CA=0.85, F1 Score = 0.85, Recall = 0.85). The differences between the integrated prediction models and Random Forest based prediction model were non-significant as observed at the fourth decimal places.

### 3.4 Evaluation of Prediction Models Based on Test Dataset

The performance of different prediction models was further evaluated on a Test Dataset of 136 experimentally characterized hexapepetides, comprising 68 aggregate forming peptides and 68 non-aggregate forming peptides. The evaluation statistics, accounting for area under ROC curve (AUC), classification accuracy (CA), F1 score, and recall metrics, for different prediction models are summarized in Table 3.

**Table 3.** A summary of benchmarking of different prediction models for identifying aggregation prone regions in protein sequences with current state of the art methods. All the methods were used in their default parameter setting to avoid any ambiguity or bias in the predictions.

| Prediction Method | Sensitivity | Specificity | F1 Score | Precision | MCC |
| --- | --- | --- | --- | --- | --- |
| APR-Score-RF | <u>0.93</u> | 0.88 | <u>0.91</u> | 0.89 | <u>0.81</u> |
| APR-Score-SVM | 0.88 | 0.88 | 0.88 | 0.88 | 0.77 |
| APR-Score-NN | 0.84 | 0.84 | 0.84 | 0.84 | 0.68 |
| APR-Score-GB | 0.84 | 0.85 | 0.84 | 0.85 | 0.69 |
| APR-Score-LR | 0.88 | 0.84 | 0.86 | 0.85 | 0.72 |
| TANGO | 0.35 | <u>1.00</u> | 0.52 | <u>1.00</u> | 0.46 |
| PASTA 2.0 | 0.21 | 0.98 | 0.34 | 0.93 | 0.31 |
| MetAmyl | 0.66 | 0.93 | 0.76 | 0.90 | 0.61 |
| ANuPP | 0.71 | 0.96 | 0.81 | 0.94 | 0.68 |
| AGGRESCAN | 0.62 | 0.95 | 0.74 | 0.93 | 0.61 |
| WALTZ | 0.24 | <u>1.00</u> | 0.38 | <u>1.00</u> | 0.36 |

It is observed that all the prediction methods mimicked their performances on the Test Dataset, similar to the performances in 10-fold cross validation. Notably, the peptides in the Test Dataset were completely exclusive of the peptides used in the Training Dataset for development of the proposed prediction models. The Random Forest based prediction model (APR-Score-RF) performed the best as compared to other prediction models in terms of area under ROC curve (AUC), classification accuracy (CA), F1 score, precision, and recall metrics. Also, the Logistic Regression based prediction model (APR-Score-LR) performed almost the same as Random Forest based prediction model (APR-Score-RF) in terms of AUC and precision metrics. Further, the performance of the prediction models was analyzed through ROC curves to identify the optimum threshold for individual model without compromising the sensitivity and specificity, as depicted in figure 2.

The ROC curves are shown in Figure 2 where the false positive rate (x-axis) and plotted against true positive rate (y-axis). The false positive rate represents the subtraction of specificity from 1 (1 - specificity) while the true positive rate represents sensitivity of the prediction models in identifying aggregation prone peptides. The threshold with optimum tradeoff between sensitivity and specificity are marked with circle in the ROC curves (Figure 1) and the corresponding metrics for sensitivity, specificity, and thresholds are summarized in the embedded table in Figure 1. It is worth mentioning that the threshold values for different prediction models can be adjusted as per the rationale of the experiment involving the protein sequences to be screened for aggregation prone regions.

**Figure 2.**
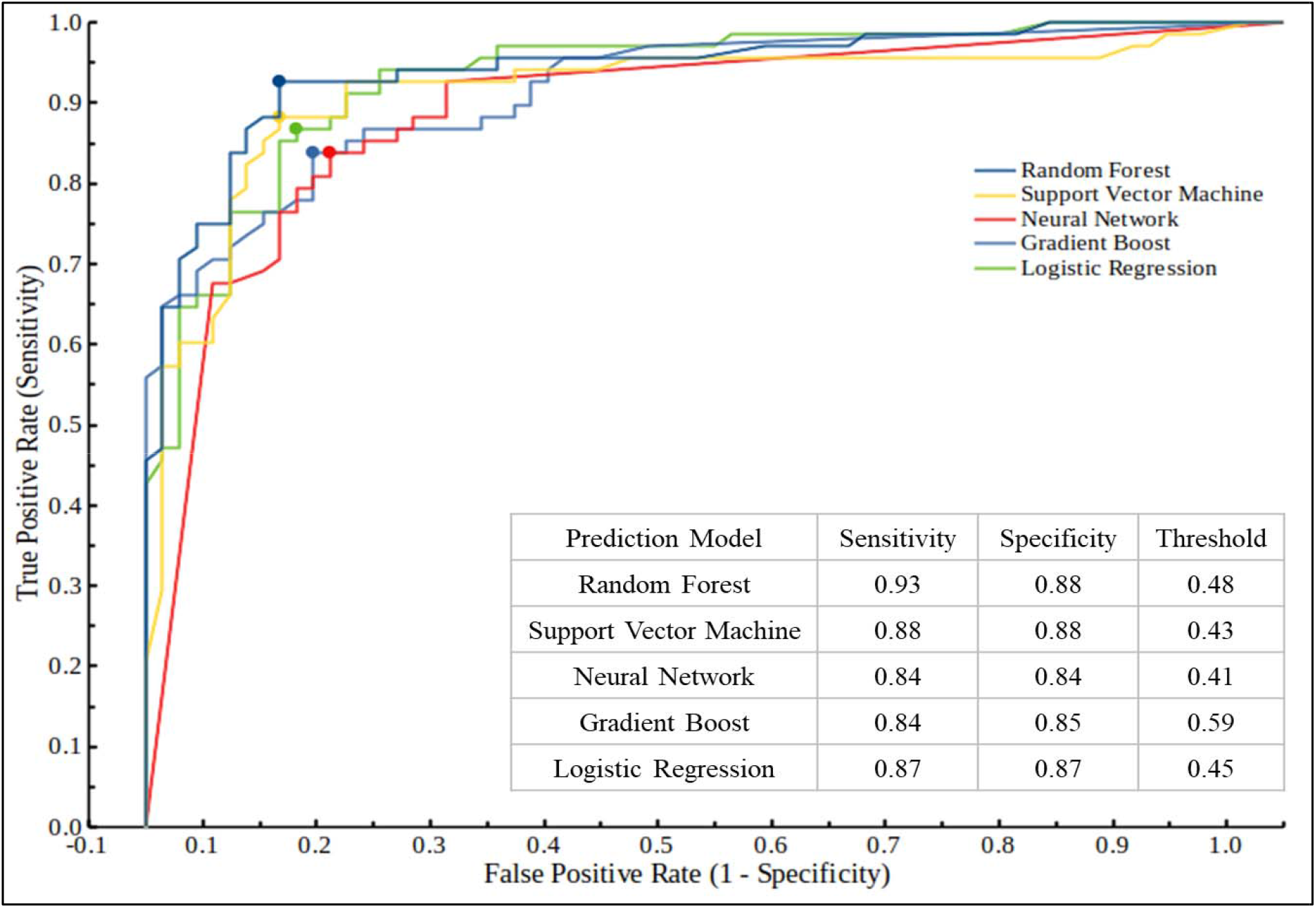
A ROC curve analysis of different prediction models on the Test Dataset. The optimum threshold values for best combination of sensitivity and specificity for individual prediction models are represented with circle. The embedded table summarizes the sensitivity and specificity at the optimum thresholds.

### 3.5 Benchmarking of Prediction Models with Current State-of-the-Art Methods

The performance of different prediction models developed in this study for identifying the aggregation prone regions in protein sequences is benchmarked with six different state-of-the-art methods, viz. TANGO (Fernandez-Escamilla, et al., 2004; Rousseau, et al., 2006), PASTA 2.0 (Walsh, et al., 2014), MetAmyl (Emily, et al., 2013), ANuPP (Prabakaran, et al., 2021), WALTZ (Beerten, et al., 2015; Louros, et al., 2020), and AGGRESCAN (Conchillo-Solé, et al., 2007; de Groot, et al., 2012). A brief description of these methods is provided in supplementary. All these methods were executed on the Test Dataset of 168 hexapeptides and their predictions were utilized for calculating different evaluation metrics, viz. sensitivity, specificity, F1 score, precision, recall, and MCC. Notably, all these methods were executed with their default parameters to avoid any type of ambiguity. It is worth mentioning that the list of state-of-the-art methods selected here for benchmarking is not exhaustive and there are several other methods that require protein tertiary structure (predicted or experimental) to perform similar predictions. Considering the overall rationale of developing this method to screen high-throughput computationally generated protein sequences, only the methods that use protein sequences as input are selected. The evaluation metrics for benchmarking of developed prediction models in this study are summarized in Table 3. Additional statistics corresponding to the predictions by different methods are provided in supplementary Table S6. The evaluation statistics for the different prediction models (APR-Scores-RF, -SVM, -NN, -GB, and -LR) are based on the threshold scores mentioned in Figure 2. The default thresholds were used for the other methods considered for benchmarking.

It is observed that the TANGO, PASTA 2.0 and WALTZ predictions showed very high specificity, i.e., the efficiency of correctly identifying non-aggregating peptides. However, the performance for identifying aggregating peptides was observed to be below par as compared to all other methods summarized in Table 3. The F1 Score metric indicates the predictive performance of individual methods by detailing their categorical performances. It takes account of accurately identified aggregating peptides (true positives), wrongly identified aggregating peptides (false positives), accurately identified non-aggregating peptides (true negatives), and wrongly identified non-aggregating peptides (false negatives). Likewise, MCC is another very informative statistical metric, equivalent to chi-square metrics for a contingency table, for evaluating the prediction models, where the higher value represents better predictive accuracy.

Among the developed prediction models, APR-Score-RF showed high MCC (0.81) and F1-Score (0.91), demonstrating its better predictive ability in distinguishing aggregating and non-aggregating regions. Among the other methods used for benchmarking, ANuPP performed reasonably well, followed by MetAmyl and AGGRESCAN. Notably, a recent version of AGGRESCAN (named AGGRESCAN-3D) that requires protein tertiary structure for identifying the aggregation prone regions, was not considered in the benchmarking, as only the methods requiring protein/peptide sequence(s) were considered. All the variants of APR-Score (-RF, -SVM, -NN, -GB, and -LR) showed improved predictions for identifying aggregation prone regions /peptides as compared to other current state of art methods, accounted for the benchmarking.

### 3.6 Implementation and Availability

The developed prediction models are made available at http://github.com/TII-BRC/MBG/tree/APR-Score. The execution of APR-Score is highly computational and time efficient. The pre-processing of the various features for a target protein sequence (length ∼ 300 aa) takes about 2 minutes on a single processor. The pre-trained prediction models can be directly used for identification of aggregation prone regions through Scikit-learn in Linux-based OS and through Orange Data Mining, an open-source machine learning tool, in Windows-based OS. The scripts for pre-processing and the pre-trained models may be executed in single sequence mode or multiple sequences (batch processing) mode. The developed prediction models may be used with command-line interface (CLI) or graphic-user interface (GUI).

## 4. Conclusion

Identifying aggregation-prone regions in protein sequences is crucial for understanding protein aggregation diseases such as Alzheimer’s, Parkinson’s, and prion diseases as well as for design and engineering novel proteins and modulating the activity of existing proteins for different rationales. The integration of multiple sequence and structure-based features and machine learning techniques, APR-Score provides a reliable and accurate means of identifying aggregation-prone regions within protein sequences. By offering robust predictive models, APR-Score equips researchers with a valuable tool to expedite the identification of potentially troublesome protein segments. This capability is instrumental in advancing our understanding of the molecular mechanisms underlying protein aggregation-related diseases and fast-tracking various protein design and engineering regimes. Furthermore, APR-Score’s predictive accuracy has been validated and benchmarked with current state of the art, cementing its practical utility in real-world applications. As our knowledge of protein aggregation continues to evolve, APR-Score remains adaptable, with the potential for continuous improvement through updated datasets and feature refinement, to unravel the complexities of protein aggregation. In essence, APR-Score not only enhances our ability to comprehend the mechanisms behind protein aggregation but also offers promise in the development of innovative strategies for efficient protein design and engineering to aid the development of targeted therapies and diagnostic tools.

## Supporting information

adopted from previous experimental studies and provided in supplementary information (Table S1)

## Authors’ contributions

RK: conceptualization, methodology, software, validation, writing – original draft. TL: methodology, Writing – review and editing. RK and TL read and approved the final manuscript.

## Acknowledgements

Rahul Kaushik and Thomas Launey are thankful to Biotechnology Research Center, Technology Innovation Institute, Abu Dhabi, UAE for its support and resources.

## References

Beerten, J., et al. WALTZ-DB: a benchmark database of amyloidogenic hexapeptides. Bioinformatics 2015;31(10):1698–1700.

Breydo, L., et al. Effects of Polymer Hydrophobicity on Protein Structure and Aggregation Kinetics in Crowded Milieu. Biochemistry 2015;54(19):2957–2966.

Burdukiewicz, M., et al. AmyloGraph: a comprehensive database of amyloid-amyloid interactions. Nucleic Acids Res 2023;51(51D):D352–d357.

Chiti, F. and Dobson, C.M. Protein misfolding, functional amyloid, and human disease. Annu Rev Biochem 2006;75:333–366.

Chiti, F. and Dobson, C.M. Amyloid formation by globular proteins under native conditions. Nat Chem Biol 2009;5(1):15–22.

Co, N.T., Li, M.S. and Krupa, P. Computational Models for the Study of Protein Aggregation. Methods Mol Biol 2022;2340:51–78.

Conchillo-Solé, O., et al. AGGRESCAN: a server for the prediction and evaluation of “hot spots” of aggregation in polypeptides. BMC Bioinformatics 2007;8:65.

Cukalevski, R., et al. Role of aromatic side chains in amyloid β-protein aggregation. ACS Chem Neurosci 2012;3(12):1008–1016.

de Groot, N.S., et al. AGGRESCAN: method, application, and perspectives for drug design. Methods Mol Biol 2012;819:199–220.

de Jong, D.H., et al. Improved Parameters for the Martini Coarse-Grained Protein Force Field. J Chem Theory Comput 2013;9(1):687–697.

Ebo, J.S., et al. Using protein engineering to understand and modulate aggregation. Curr Opin Struct Biol 2020;60:157–166.

Emily, M., Talvas, A. and Delamarche, C. MetAmyl: a METa-predictor for AMYLoid proteins. PLoS One 2013;8(11):e79722.

Eshari, F., et al. Prediction of protein aggregation propensity employing SqFt-based logistic regression model. Int J Biol Macromol 2023;249:126036.

Fang, Y., et al. Identification of properties important to protein aggregation using feature selection. BMC Bioinformatics 2013;14:314.

Fernandez-Escamilla, A.M., et al. Prediction of sequence-dependent and mutational effects on the aggregation of peptides and proteins. Nat Biotechnol 2004;22(10):1302–1306.

Gallardo, J., Escalona-Noguero, C. and Sot, B. Role of α-Synuclein Regions in Nucleation and Elongation of Amyloid Fiber Assembly. ACS Chem Neurosci 2020;11(6):872–879.

Galzitskaya, O.V., Garbuzynskiy, S.O. and Lobanov, M.Y. Prediction of amyloidogenic and disordered regions in protein chains. PLoS Comput Biol 2006;2(12):e177.

Garbuzynskiy, S.O., Lobanov, M.Y. and Galzitskaya, O.V. FoldAmyloid: a method of prediction of amyloidogenic regions from protein sequence. Bioinformatics 2010;26(3):326–332.

Guo, S. and Akhremitchev, B.B. Packing density and structural heterogeneity of insulin amyloid fibrils measured by AFM nanoindentation. Biomacromolecules 2006;7(5):1630–1636.

Guy, H.R. Amino acid side-chain partition energies and distribution of residues in soluble proteins. Biophys J 1985;47(1):61–70.

Hartl, F.U. Protein Misfolding Diseases. Annu Rev Biochem 2017;86:21–26.

Houben, B., Rousseau, F. and Schymkowitz, J. Protein structure and aggregation: a marriage of necessity ruled by aggregation gatekeepers. Trends Biochem Sci 2022;47(3):194–205.

Housmans, J.A.J., et al. A guide to studying protein aggregation. Febs j 2023;290(3):554–583.

Jones, S., Marin, A. and Thornton, J.M. Protein domain interfaces: characterization and comparison with oligomeric protein interfaces. Protein Eng 2000;13(2):77–82.

Kaushik, R. and Zhang, K.Y.J. A protein sequence fitness function for identifying natural and nonnatural proteins. Proteins 2020;88(10):1271–1284.

Kaushik, R. and Zhang, K.Y.J. An integrated protein structure fitness scoring approach for identifying native-like model structures. Comput Struct Biotechnol J 2022;20:6467–6472.

Kaushik, R. and Zhang, K.Y.J. ProFitFun: a protein tertiary structure fitness function for quantifying the accuracies of model structures. Bioinformatics 2022;38(2):369–376.

Knowles, T.P., Vendruscolo, M. and Dobson, C.M. The amyloid state and its association with protein misfolding diseases. Nat Rev Mol Cell Biol 2014;15(6):384–396.

Langenberg, T., et al. Thermodynamic and Evolutionary Coupling between the Native and Amyloid State of Globular Proteins. Cell Rep 2020;31(2):107512.

Louros, N., et al. WALTZ-DB 2.0: an updated database containing structural information of experimentally determined amyloid-forming peptides. Nucleic Acids Res 2020;48(48D):D389–d393.

Madhusudan Makwana, K. and Mahalakshmi, R. Implications of aromatic-aromatic interactions: From protein structures to peptide models. Protein Sci 2015;24(12):1920–1933.

Maurer-Stroh, S., et al. Exploring the sequence determinants of amyloid structure using position-specific scoring matrices. Nat Methods 2010;7(3):237–242.

Maurya, G.P., et al. Hydrophobicity Directed Chiral Self-Assembly and Aggregation-Induced Emission: Diacetylene-Cored Pseudopeptide Chiral Dopants. Angew Chem Int Ed Engl 2022;61(42):e202209806.

Meric, G., et al. Challenges for design of aggregation-resistant variants of granulocyte colony-stimulating factor. Biophys Chem 2021;277:106630.

Naderi-Manesh, H., et al. Prediction of protein surface accessibility with information theory. Proteins 2001;42(4):452–459.

O’Donnell, C.W., et al. A method for probing the mutational landscape of amyloid structure. Bioinformatics 2011;27(13):i34–42.

Ono, K. and Watanabe-Nakayama, T. Aggregation and structure of amyloid β-protein. Neurochem Int 2021;151:105208.

Orlando, G., et al. Accurate prediction of protein beta-aggregation with generalized statistical potentials. Bioinformatics 2020;36(7):2076–2081.

Peng, C., Trojanowski, J.Q. and Lee, V.M. Protein transmission in neurodegenerative disease. Nat Rev Neurol 2020;16(4):199–212.

Prabakaran, R., et al. ANuPP: A Versatile Tool to Predict Aggregation Nucleating Regions in Peptides and Proteins. J Mol Biol 2021;433(11):166707.

Ptak-Kaczor, M., et al. Solubility and Aggregation of Selected Proteins Interpreted on the Basis of Hydrophobicity Distribution. Int J Mol Sci 2021;22(9).

Rawat, P., et al. CPAD 2.0: a repository of curated experimental data on aggregating proteins and peptides. Amyloid 2020;27(2):128–133.

Rousseau, F., Schymkowitz, J. and Serrano, L. Protein aggregation and amyloidosis: confusion of the kinds? Curr Opin Struct Biol 2006;16(1):118–126.

Santos, J., et al. Computational prediction of protein aggregation: Advances in proteomics, conformation-specific algorithms and biotechnological applications. Comput Struct Biotechnol J 2020;18:1403–1413.

Seuma, M., Lehner, B. and Bolognesi, B. An atlas of amyloid aggregation: the impact of substitutions, insertions, deletions and truncations on amyloid beta fibril nucleation. Nat Commun 2022;13(1):7084.

Shaytan, A.K., Shaitan, K.V. and Khokhlov, A.R. Solvent accessible surface area of amino acid residues in globular proteins: correlation of apparent transfer free energies with experimental hydrophobicity scales. Biomacromolecules 2009;10(5):1224–1237.

Sigurdson, C.J., Bartz, J.C. and Glatzel, M. Cellular and Molecular Mechanisms of Prion Disease. Annu Rev Pathol 2019;14:497–516.

Skibiszewska, S., et al. Influence of short peptides with aromatic amino acid residues on aggregation properties of serum amyloid A and its fragments. Arch Biochem Biophys 2020;681:108264.

Smith, D.J. and Shell, M.S. Can Simple Interaction Models Explain Sequence-Dependent Effects in Peptide Homodimerization? J Phys Chem B 2017;121(24):5928–5943.

Stankovic, I.M., et al. Role of aromatic amino acids in amyloid self-assembly. Int J Biol Macromol 2020;156:949–959.

Teng, P.K. and Eisenberg, D. Short protein segments can drive a non-fibrillizing protein into the amyloid state. Protein Eng Des Sel 2009;22(8):531–536.

Varadi, M., et al. AmyPro: a database of proteins with validated amyloidogenic regions. Nucleic Acids Res 2018;46(46D):D387–d392.

Vendruscolo, M. and Fuxreiter, M. Towards sequence-based principles for protein phase separation predictions. Curr Opin Chem Biol 2023;75:102317.

Walsh, I., et al. PASTA 2.0: an improved server for protein aggregation prediction. Nucleic Acids Res 2014;42(Web Server issue):W301–307.

Xiao, H., et al. Static and dynamic disorder in Aβ40 fibrils. Biochem Biophys Res Commun 2022;610:107–112.

Zbilut, J.P., et al. Entropic criteria for protein folding derived from recurrences: six residues patch as the basic protein word. FEBS Lett 2006;580(20):4861–4864.

Zhang, Y., et al. An improved profile-level domain linker propensity index for protein domain boundary prediction. Protein Pept Lett 2011;18(1):7–16.

Zhang, Z., Chen, H. and Lai, L. Identification of amyloid fibril-forming segments based on structure and residue-based statistical potential. Bioinformatics 2007;23(17):2218–2225.

