## Supplementary material for "Decoding Protein Aggregation through Computational Approach: Identification and Scoring of Aggregation-Prone Regions in Protein Sequences": adopted from previous experimental studies and provided in supplementary information (Table S1)

***Supplementary Tables***

**Table S1.** The normalized scores for hydrophobicity (base scores are adopted Naderi-Manesh Scale) where positive values represent hydrophobic nature of residues and vice-versa, packing density (base scores are adopted from Galzitskaya et al., 2006) where positive values show densely packed residues and vice-versa, linker index (base scores adopted from Bae et al., 2005) where positive values indicate highly linked (interacting) residues and vice-versa, mean polarity (base scores are adopted Naderi-Manesh Scale) where positive scores represent non-polar residues and vice-versa, and partition energies (Guy, 1985).

| Amino Acid | Hydrophobicity | Packing Density | Linker Index | Mean Polarity | Partition Energy |
| --- | --- | --- | --- | --- | --- |
| Alanine (A) | 0.3723 | -0.3713 | -0.0705 | -0.1221 | -0.6246 |
| Arginine (R) | -0.7024 | -0.1325 | -0.2686 | -0.7228 | 0.2258 |
| Asparagine (N) | -0.4098 | -0.6868 | -0.2737 | -0.4455 | 0.2977 |
| Aspartate (D) | -0.3805 | -0.9527 | -0.3738 | -0.6920 | 0.3320 |
| Cysteine (C) | 1.0000 | 0.3432 | 1.0000 | 1.0000 | 0.3168 |
| Glutamine (Q) | -0.6244 | -0.5471 | -0.3303 | -0.6380 | 1.0000 |
| Glutamate (E) | -0.5610 | -0.9437 | -0.4889 | -0.6688 | -0.9659 |
| Glycine (G) | -0.0634 | -1.0000 | -0.2003 | -0.3916 | -0.1760 |
| Histidine (H) | -0.2683 | 0.0032 | 0.2194 | -0.3103 | -0.4832 |
| Isoleucine (I) | 0.7737 | 0.6282 | 0.4326 | 0.9604 | 0.7773 |
| Leucine (L) | 0.7518 | 0.5660 | 0.3877 | 0.8811 | -0.7075 |
| Lysine (K) | -1.0000 | -0.8783 | -0.5615 | -0.9846 | -0.5076 |
| Methionine (M) | 0.5328 | 0.4604 | -0.0639 | 0.9287 | 0.2121 |
| Phenylalanine (F) | 0.7883 | 0.8394 | 0.5862 | 0.9287 | 0.1607 |
| Proline (P) | -0.3854 | -0.9211 | -1.0000 | -1.0000 | 0.3731 |
| Serine (S) | -0.1268 | -0.7664 | -0.4537 | -0.4609 | 0.0716 |
| Threonine (T) | -0.0146 | -0.3849 | -0.2556 | -0.2838 | -0.5514 |
| Tryptophan (W) | 0.5036 | 1.0000 | 0.6389 | 0.6195 | -0.4588 |
| Tyrosine (Y) | 0.0803 | 0.6586 | 0.3833 | 0.1835 | 0.0476 |
| Valine (V) | 0.7883 | 0.3519 | 0.2459 | 0.7860 | -1.0000 |

**Table S2.** A summary of Shannon’s entropy at different position specificity of amino acid residues in hexapeptides.

| Amino Acid | Position 1 | Position 2 | Position 3 | Position 4 | Position 5 | Position 6 |
| --- | --- | --- | --- | --- | --- | --- |
| Alanine (A) | 0.342 | 0.413 | 0.142 | 0.530 | 0.179 | 0.453 |
| Arginine (R) | 0.189 | 0.062 | 0.000 | 0.103 | 0.000 | 0.113 |
| Asparagine (N) | 0.629 | 0.413 | 0.454 | 0.735 | 0.515 | 0.519 |
| Aspartate (D) | 0.223 | 0.221 | 0.059 | 0.000 | 0.000 | 0.324 |
| Cysteine (C) | 0.063 | 0.062 | 0.142 | 0.143 | 0.059 | 0.156 |
| Glutamine (Q) | 0.439 | 0.479 | 0.392 | 0.143 | 0.439 | 0.559 |
| Glutamate (E) | 0.151 | 0.312 | 0.269 | 0.242 | 0.104 | 1.000 |
| Glycine (G) | 0.674 | 0.312 | 0.346 | 0.270 | 0.497 | 0.324 |
| Histidine (H) | 0.417 | 0.187 | 0.177 | 0.103 | 0.179 | 0.000 |
| Isoleucine (I) | 0.462 | 0.760 | 0.370 | 1.000 | 1.000 | 0.353 |
| Leucine (L) | 0.503 | 0.591 | 0.434 | 0.701 | 0.417 | 0.430 |
| Lysine (K) | 0.368 | 0.187 | 0.059 | 0.143 | 0.000 | 0.264 |
| Methionine (M) | 0.286 | 0.187 | 0.210 | 0.000 | 0.144 | 0.353 |
| Phenylalanine (F) | 0.314 | 0.458 | 0.876 | 0.436 | 0.597 | 0.559 |
| Proline (P) | 0.000 | 0.062 | 0.000 | 0.000 | 0.104 | 0.000 |
| Serine (S) | 1.000 | 0.413 | 0.322 | 0.297 | 0.271 | 0.632 |
| Threonine (T) | 0.189 | 1.000 | 0.210 | 0.394 | 0.212 | 0.648 |
| Tryptophan (W) | 0.223 | 0.000 | 0.295 | 0.323 | 0.243 | 0.194 |
| Tyrosine (Y) | 0.368 | 0.537 | 0.142 | 0.348 | 0.497 | 0.379 |
| Valine (V) | 0.596 | 0.499 | 1.000 | 0.270 | 0.373 | 0.519 |

**Table S3**. A summary of Principal Component Analysis for selection of non-redundant parameter to capture 99.9% cumulative variance from the compiled parameters (n =22) in the Training Dataset of hexapeptides. The extent of information gain ratio, and ANOVA are used as four parameters (S.N. 19-22) were identified as redundant and thus discarded while developing the prediction models.

| S. N. | Parameter | Gain ratio | ANOVA |
| --- | --- | --- | --- |
| 1 | Overall Packing Density Score of Hexapeptide | 0.195 | 430.79 |
| 2 | Ratio of Densely to Loosely Packed Residues | 0.192 | 404.36 |
| 3 | Average Number of Densely Packed Residues | 0.143 | 262.41 |
| 4 | Average Number of Loosely Packed Residues | 0.126 | 259.68 |
| 5 | Overall Mean Polarity Score of Hexapeptide | 0.188 | 421.47 |
| 6 | Ratio of Polar to Non-Polar Residues | 0.174 | 398.10 |
| 7 | Average Number of Non-Polar Residues | 0.152 | 262.59 |
| 8 | Average Number of Polar Residues | 0.076 | 114.23 |
| 9 | Overall Linker Index Score of Hexapeptide | 0.172 | 369.62 |
| 10 | Ratio of Linker to Non-Linker Residues | 0.164 | 367.56 |
| 11 | Average Number of Linker Residues | 0.136 | 226.99 |
| 12 | Average Number of Non-Linker Residues | 0.110 | 203.12 |
| 13 | Overall Hydrophobicity Score of Hexapeptide | 0.146 | 296.03 |
| 14 | Ratio of Hydrophobic to Non-Hydrophobic Residues | 0.123 | 275.31 |
| 15 | Average Number of Hydrophobic Residues | 0.123 | 243.08 |
| 16 | Average Number of Non-Hydrophobic Residues | 0.066 | 78.26 |
| 17 | Overall Sequence-Structure Compatibility (CSS) Score | 0.085 | 58.07 |
| 18 | Overall Shannon’s Entropy Score of Hexapeptide | 0.028 | 40.57 |
| 19 | Average Negative Partition Energy | 0.019 | 33.48 |
| 20 | Ratio of Positive to Negative Partition Energy Residues | 0.017 | 21.12 |
| 21 | Overall Partition Energy of Hexapeptide | 0.012 | 20.54 |
| 22 | Average Positive Partition Energy | 0.008 | 3.62 |

**Table S4.** A summary of sensitivity, specificity, precision, true positive rate, and false positive rate analysis (at threshold p = 0.50) of different machine learning based prediction models.

| Evaluation Metric | APR-Score-RF | APR-Score-SVM | APR-Score-NN | APR-Score-GB | APR-Score-LR |
| --- | --- | --- | --- | --- | --- |
| True Positive Rate | 0.872 | 0.864 | 0.821 | 0.835 | 0.835 |
| False Positive Rate | 0.176 | 0.194 | 0.209 | 0.165 | 0.194 |

**Table S5.** A summary of performance evaluation of different machine learning based prediction models in terms of area under ROC curve (AUC), classification accuracy (CA), F1 score, precision, and recall metrics to identify the experimentally characterized aggregation prone peptides.

| Prediction Model | AUC | CA | F1 Score | Precision | Recall |
| --- | --- | --- | --- | --- | --- |
| APR-RF | 0.94 | 0.89 | 0.89 | 0.88 | 0.90 |
| APR-SVM | 0.91 | 0.88 | 0.87 | 0.88 | 0.87 |
| APR-NN | 0.91 | 0.83 | 0.83 | 0.84 | 0.82 |
| APR-GB | 0.92 | 0.84 | 0.84 | 0.84 | 0.84 |
| APR-LR | 0.93 | 0.85 | 0.84 | 0.87 | 0.81 |

**Table S6.** Additional statistics for benchmarking of different prediction models for identifying aggregation prone regions in protein sequences with current state of the art methods. All the methods were used in their default parameter setting to avoid any ambiguity or bias in the predictions.

| Method | True Positive | False Negative | True Negative | False Positive |
| --- | --- | --- | --- | --- |
| APR-RF | 63 | 5 | 60 | 8 |
| APR-SVM | 60 | 8 | 60 | 8 |
| APR-NN | 57 | 11 | 57 | 11 |
| APR-GB | 57 | 11 | 58 | 10 |
| APR-LR | 60 | 8 | 57 | 11 |
| TANGO | 24 | 44 | 68 | 0 |
| PASTA 2.0 | 14 | 54 | 67 | 1 |
| MetAmyl | 45 | 23 | 63 | 5 |
| ANuPP | 48 | 20 | 65 | 3 |
| AggreScan | 42 | 26 | 65 | 3 |
| WALTZ | 16 | 52 | 68 | 0 |

***Supplementary Note - I***

A brief description of methods used for benchmarking the performance of different prediction models for identifying aggregation prone regions in protein sequences.

(a) TANGO: The TANGO method predicts the cross-beta aggregating segments in the peptide /protein sequences by calculating a Boltzmann distribution-based partition function that implementing information and propensities derived from four different secondary structural states, viz. α-helices, β-turns, α-helical aggregations, and β-strand aggregations. The TANGO method is available as standalone as well as webserver. It predicts percentage of aggregation and helical aggregation at residue and peptide/ protein sequence levels. Further information regarding various parameter used in TANGO can be accessed at <http://tango.crg.es>.
(b) PASTA (2.0): It predicts protein aggregation propensities for protein sequence by utilizing a diverse set of features derived from aggregating peptides /proteins. The PASTA 2.0 energy function accounts for the stability of potential cross-beta pairings. In addition to predicting the aggregation prone regions, PASTA 2.0 also delivers information about potential intrinsic disordered regions and different secondary structural elements for the input protein sequences. The PASTA 2.0 method is freely available as standalone as well as webserver for academic users and on request for non-academic users. The PASTA 2.0 server also supports batch prediction and can be accessed at <http://protein.bio.unipd.it/pasta2/>.
(c) MetAmyl: It is a meta predictor for identifying aggregation regions (amyloid hot spots) from the protein sequences. It integrates different existing methods for predicting aggregation prone regions by implementing logistic regression model. The different methods integrated in MetAmyl included PASTA, SALSA, TANGO, PAFIG, AGGRESCAN, Waltz, and AmylFold. The weighted combination of the 11 scores derived from the integrating methods showed an improved performance over their individual efficiencies. The MetAmyl is available as webserver and supports batch predictions for multiple peptide /protein sequences (accessible at <http://metamyl.genouest.org/e107_plugins/metamyl_aggregation/db_prediction_meta.php>).
(d) ANuPP: This method performs prediction of aggregation nucleating regions in peptides and protein sequences. The ANuPP is a web-based meta-classifier method that implements atomic-level features as ensemble classifier to explore aggregation nucleation in peptides and proteins. The method is available as webserver, available at <https://web.iitm.ac.in/bioinfo2/ANuPP>, and supports batch processing for multiple peptide /protein sequences.
(e) WALTZ: It is a Position Specific Substitution Matrices (PSSM) based statistical methods derived from the physicochemical properties of aggregating (amyloid forming) hexapeptides along with structural properties. The Waltz method identifies amyloid sequences / amyloid forming regions and beta-sheet aggregates. The amino acid residue specific PSSM scores are log-odd scores derived from amyloid and non-amyloid forming hexapeptides. The final scoring function utilized 19 physicochemical properties, screened statistically to extract maximum information from the dataset of amyloid and non-amyloid forming hexapeptides. Waltz is available as webserver at <https://waltz.switchlab.org/> and supports batch processing of multiple peptide/protein sequences.
(f) AGGRESCAN: It identifies aggregation-prone regions in protein sequences by implementing an amino acid residues level aggregation propensity scale derived from the experimental dataset of short and specific protein sequence segments that were known to modulate aggregation in proteins. In addition to aggregation-prone region identification, it can also perform mutational analysis on protein aggregation and aggregation features comparison in a batch mode for multiple protein sequences. The webserver for AGGRESCAN can be accessed at <http://bioinf.uab.es/aggrescan/>.
